# Surface curvature promotes coordinated extracellular matrix remodeling and mechanotransductive signaling during osteoblast adaptation

**DOI:** 10.64898/2026.10.27.762219

**Authors:** Heidi Pertl-Obermeyer, Angelika Lahnsteiner, Barbara Schamberger, Gerhard Obermeyer, John W. C. Dunlop

## Abstract

**Background:** Substrate curvature is an important biophysical cue regulating cell behavior through mechanotransduction. However, how three-dimensional curvature regulates osteoblast transcriptional adaptation remains poorly understood. We investigated the temporal transcriptional response of pre-osteoblastic cells to defined substrate curvature.

**Methods:** Murine MC3T3-E1 pre-osteoblasts were cultured on three-dimensional polydimethylsiloxane capillary bridges or flat substrates for 7 and 23 days to investigate the effects of surface curvature on gene expression. A genome-wide transcriptomic approach was employed to comprehensively characterize differential gene expression, followed by a focused analysis of selected biological pathways, including osteogenic signaling, extracellular matrix (ECM) organization, cell–matrix adhesion, cytoskeletal regulation, and mechanosensitive signaling.

**Results:** Substrate curvature induced pronounced, time-dependent transcriptional changes. At day 7, curved substrates activated immediate-early and stress-responsive genes, including *Egr1, Egr2, Fos, Junb,* and *Nr4a1*. By day 23, the response shifted toward ECM remodeling, receptor-mediated signaling, and structural adaptation, including increased expression of *Ankrd1, Ereg, Serpine1,* and *Plaur*. Despite these changes, canonical osteogenic transcription factors, including *Runx2, Cb@, Osx/Sp7,* and *Atf4*, showed limited alterations. In contrast, BMP, WNT, FGF, and Notch signaling, ECM components, integrins, and actin-associated genes were dynamically regulated, whereas *Piezo1, Piezo2,* and most septins remained comparatively stable.

**Conclusions:** Substrate curvature regulates osteoblast transcriptional adaptation primarily through coordinated changes in ECM organization, cell–matrix interactions, cytoskeletal architecture, and signaling networks rather than reprogramming the canonical osteogenic transcriptional machinery.

## Introduction

The curvature of the cell membrane is a fundamental physical element that can regulate gene expression and cellular behavior through mechanobiological processes (Assoian et al. 2019, Schamberger et al. 2023, Luciano et al. 2024, Arnold et al. 2025). Single cells detect the (surface) geometry of their environment through interconnected mechanosensory systems that couple extracellular matrix (ECM) interactions with intracellular force transmission. Therefore, different surface curvatures will alter the cell membrane geometry and the tension of the adherent cytoskeleton resulting in changes in focal adhesion and actomyosin contractility. The transduction of mechanical signals into biochemical activities requires so called mechanotransduction signaling pathways, which involve molecular processes at the cell surface as well as the nucleus, being mechanically associated with the ECM through the cytoskeleton (Wang 2017, Werner et al. 2017). In addition to chemical components of the ECM, which affect cell proliferation, migration and cell differentiation (Assoian et al. 2019, Callens et al. 2020), the surface geometry itself, e.g. curvature, influences single cells (Werner et al. 2017, Callens et al. 2020, Arnold et al. 2025). Pieuchot et al. (2018) demonstrated that adherent osteoblasts actively sense cell-scale curvature landscapes and preferentially migrate according to local geometric features, a phenomenon called *curvotaxis*. These findings established that substrate curvature is an important determinant of cell behavior and that geometry is a key regulator of mechanotransduction (Pieuchot et al. 2018). Furthermore, these single cell studies clearly indicate that curvature sensing is driven by mechano-signaling due to localized changes in the cell-membrane curvature. This requires a characteristic length scale of curvature that is much smaller than that of the single cell. It is thus likely that when this length scale reaches a larger than that of a cell, curvature sensing will stop. Surprisingly, many recent studies have shown that when cells become confluent and a tissue is formed, small surface curvatures modulate tissue growth rates (Rumpler et al. 2008, Bidan et al. 2012, Bidan et al. 2013) and patterning (Bidan et al. 2016, Ehrig et al. 2019, Callens et al. 2023, Schamberger et al. 2024). Some indirect evidence, by changing cell contractility with Blebbistatin or TGF-ß suggest that this is also driven by mechanotransduction but the underlying molecular mechanisms by which many cells collectively sense and adapt to curvature remain incomplete.

The ECM, defined as the non-cellular component of tissues, plays a crucial role in curvature sensing, contributes to the organization of cells into tissues, guides cell polarization and, in addition provides and relays mechanical and biochemical signals (Naba 2024). Multiple matrix proteins and glycans form the primary components of the ECM to provide the necessary support to cells and tissue (Frantz et al. 2010). These proteins are categorized as either structural, e.g. collagens and elastin, or non-structural, like fibronectin and laminins. Other important ECM components include integrins, growth factors (GFs), and matrix metalloproteinases (MMPs, (TaipaleandKeski-Oja 1997, Kular et al. 2014)). Collagens within the ECM provide tensile strength and play a role in cell adhesion and migration. They are the predominant form of structural proteins found within the ECM, arranged into fibrils to provide the required structural integrity for the tissue. Elastin is strongly associated with collagen and plays a significant role in matrix elasticity, whereas fibronectins (FN) bind to collagens and heparan sulphate proteoglycans (HSPGs, (Kim et al. 2011)) and are involved in cell adhesion. FN is embedded into a mesh of collagen fibrils and is linked to cell surface receptors, e.g. integrins. Integrins are heterodimeric transmembrane receptors that can bind to a broad panel of different ECM molecules (Alam et al. 2007). For instance, laminins, another type of ECM glycoproteins, also bind to integrins and play a vital part in differentiation and migration, cell adhesion and are partly responsible for providing the tensile strength of the tissue (Kular et al. 2014). Therefore, any cell-specific expression of different integrins and integrin binding molecules may determine the downstream cellular responses to mechanical ECM properties, e.g. stiffness (Kim et al. 2011).

The ECM also contains a reservoir of growth factors, including insulin-like growth factors (IGFs), fibroblast growth factors (FGFs), transforming growth factor-ß (TGF-ß), and hepatocyte growth factor (HGF) that are associated with ECM proteins (Taipale and Keski-Oja 1997, Alam et al. 2007). This storage of large quantities of growth factors in the ECM as ready-to-go signaling molecules allows rapid responses to extra- and/or intercellular signals. Cells can interpret the GFs signals present in their surrounding matrix. In turn, GFs can modulate the synthesis and degradation of the ECM, and this feedback response plays a key role in tissue formation. MMPs are enzymes involved in continuous remodeling of the ECM (Kular et al. 2014) by degradation of existing and synthesis of new ECM components, thus playing an essential role in hormone-dependent tissue remodeling and tissue repair (Bosman and Stamenkovic 2003) because MMP expression itself is rapidly regulated by GFs and cytokines that participate in mechanisms to remodel ECM proteins and mechanotransductive signaling. In a simplified view, mechanical stimuli from the extracellular matrix are transmitted through integrin-containing focal adhesions to the actin cytoskeleton (Geiger et al. 2009, Hoffman et al. 2011) where they are converted into biochemical signals that regulate a variety of mechanosensitive pathways, including FAK/MAPK/ERK (Stewart et al. 2020, CrozetandLevayer 2023), BMP, WNT/β-catenin, and Notch signaling (Thompson et al. 2012), which finally control osteoblast proliferation, differentiation and as a feedback response, the extracellular matrix remodeling. Recent studies have further identified PIEZO channels as mechanically activated ion channels in bone cells (Xu et al. 2021) and septin complexes as membrane curvature sensors capable of organizing cortical cytoskeletal architecture (Shi et al. 2023). Despite the progress in the last years, it remains unclear how these mechanoresponsive components interact during cellular adaptation to curved substrates, with characteristic length scales much larger than that of a single cell.

To obtain an extensive view of processes involved in curvature-dependent sensing and signal transduction, the transcriptome of pre-osteoblastic MC3T3-E1 cells cultured as a tissue on curved surfaces in comparison with the transcriptome on flat surface-grown cells was analyzed with particular emphasis on extracellular matrix remodeling, mechanotransduction, cytoskeletal organization, and osteogenic signaling networks.

## Materials and methods

### Fabrication of capillary bridges

Capillary bridges (CBs) were fabricated as described previously (Ehrig et al. 2019, Schamberger et al. 2024). In brief, polydimethylsiloxane (PDMS) was prepared by mixing (weight plus weight) 10 parts of base agent with 1 part of curing agent of the Sylgard 184 (Dow Corning, USA). After degassing for 20 min, appropriate volumes of PDMS were transferred to cylindrical aluminium pillars with a diameter of 2 mm facing each other at a distance of 1.25 mm to produce CBs of 1.25 mm length, 2 mm end diameter and neck diameter between 1.08 – 1.18 mm. PDMS capillary bridges were cured at 120 °C for 20 min. The capillary bridges all had mean curvatures ranging from −0.066 to −0.063 mm^−1^ (calculated from the neck radius and the equations of (Gillette and Dyson 1971). These give a characteristic length of ∼15mm which is much larger than that of a single cell.

Before cell seeding, the surfaces of CBs were functionalized by O_2_ plasma treatment (K1050X, Quorum Technologies, UK with settings 60 s, 50 W) followed by an incubation in 3 % v/v aminopropyltriethoxysilane, (APTES, Sigma-Merck, Darmstadt, Germany) for 15 min at room temperature (RT) and rinsed 3x with ddH_2_O. After rinsing CBs were incubated in 1 % v/v glutaraldehyde (Carl Roth, Karlsruhe, Germany) for 30 min at RT and washed 3x with ddH_2_O. Finally, CBs were incubated with 25 µg/ml fibronectin (Sigma-Merck, Darmstadt, Germany) in phosphate-buffered saline (PBS, Sigma-Merck, Darmstadt, Germany) for 1 h at RT and washed extensively with PBS.

### Cell culture

Murine pre-osteoblastic MC3T3-E1 cells (ACC 210, DSMZ-German Collection of Microorganisms and Cell Cultures GmbH, Braunschweig, Germany,(Kodama et al. 1981)) were provided by the Ludwig Boltzmann Institute of Osteology (Vienna, Austria) and used for all experiments. Cell cultures were tested regularly for mycoplasma contamination by standard PCR method.

MC3T3-E1 were either seeded onto the surface of capillary bridges (CB) or onto a flat Petri dish (FT) at densities of 10^5^ cells/cm² and 10^4^ cells/cm^2^, respectively, and cultured in MEM (Sigma-Merck, Darmstadt, Germany) supplemented with 4,500 mg/l D-(+)-glucose (Sigma-Merck), 10% fetal bovine serum (Gibco, Life Technologies, Paisley, UK), 50 µg/ml L-ascorbic acid (Sigma-Merck), and 1% penicillin-streptomycin (Sigma-Merck). Cells on CBs or in Petri dishes (FT) were incubated at 37°C in a humified atmosphere with 5% CO_2_, with regular change of the culture medium every 2-3 days during the experiment (Ehrig et al. 2019,Schamberger et al. 2024). Cultures on CBs were performed with three biological replicates for each experiment and point in time (day 7 and day 23) using 10 CBs each, and three biological replicates for FT for each experiment and point in time (day 7 and day 23) with each sample containing cells of one Petri dish (diameter 6 cm).

### RNA extraction, sequencing and data analysis

Cells on a curved surface and in flat petri dishes were harvested at day 7 and day 23 by washing cells with PBS, followed by incubation in 0.25% trypsin-EDTA solution (Sigma-Merck) for 5 min at 37°C to detach the cells from the CBs and petri dishes (Fig. 1A). Cells from 10 CBs of each experiment and day were pooled and transferred into sterile 1.5 ml microtubes and centrifuged at 1,500 x g for 5 min. Supernatants were discarded and cells washed again in PBS by centrifugation for three times. The resulting final cell pellets were carefully resuspended in 50 µl PBS and shock frozen in liquid nitrogen. Samples from FTs were treated similarly. All samples were stored at −80°C.

**Figure 1.**
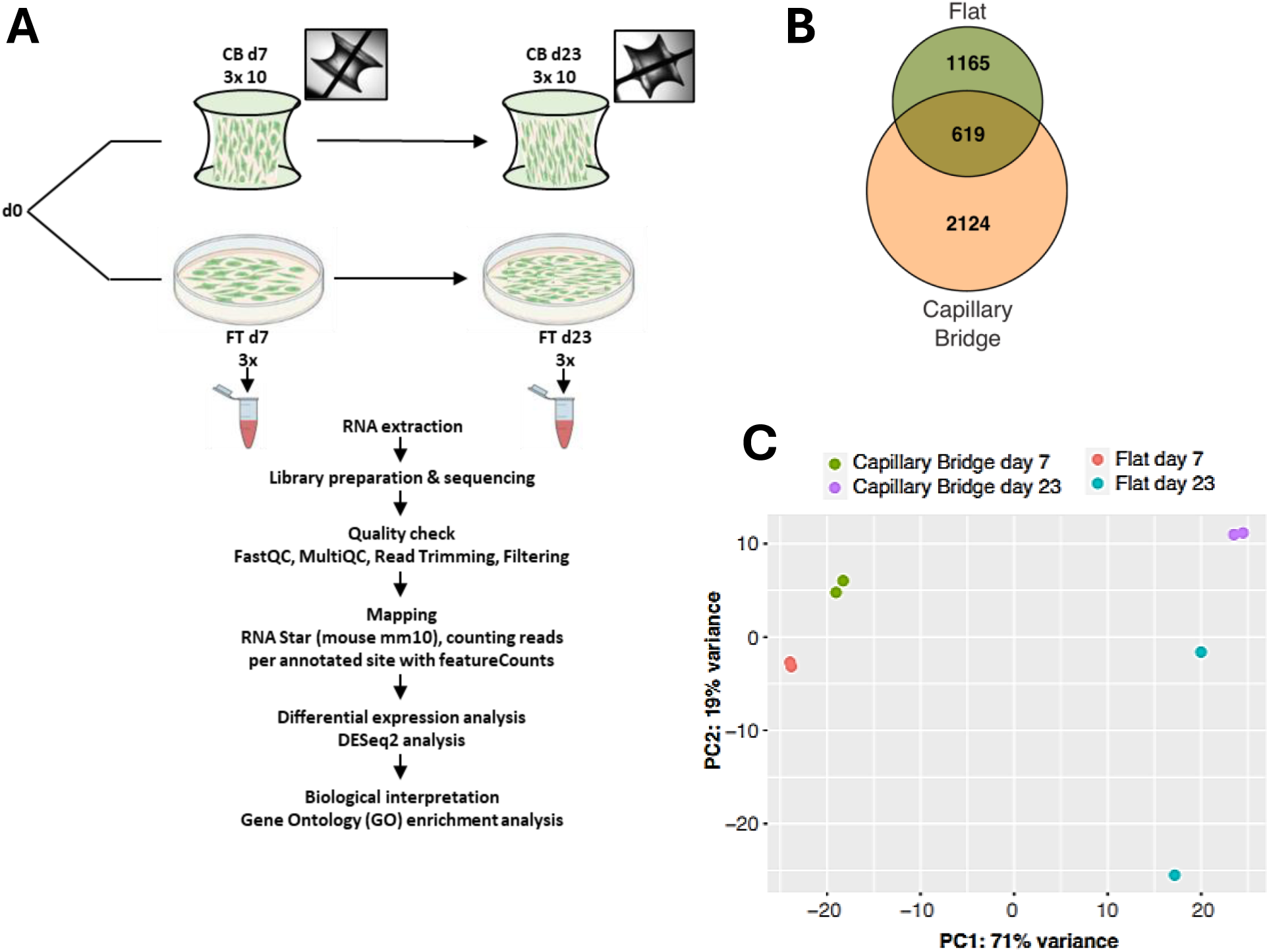
Differential gene expression of MC3T3-E1 cells on curved and flat surfaces. **A**. Schematic illustration of the experimental design and analysis workflow. The same batch of pre-osteoblastic cells were incubated either on capillary bridges (CB, curved) or in Petri dishes (FT, flat) and samples were taken after 7 and 23 days for RNA extraction to analyse the differences in gene expression. **B.** Venn diagram shows the numbers of significant differentially expressed genes at day 23 that are unique for flat-cultured (1,165) and curved-cultured (2,124) cells. 619 genes are expressed under both conditions. **C.** Principal component analysis (PCA) shows clustering of data based on the different surface geometries (CB and FT) and incubation time (day 7 and day 23).

Total RNA was extracted using the Direct-zol RNA miniprep kit (Zymo Research, Freiburg, Germany) according to the manufacturer’s guidelines. Library preparation and sequencing was conducted by Azenta GENEWIZ (Leipzig, Germany). After quality control and adapter trimming of raw fastq files, reads were aligned to the mm10 mouse reference genome using the STAR aligner (Dobin et al. 2013) tool. Only reads with a minimal mapping quality for unique mappers of 60 were used for alignment leading to approximately 50 million paired-end reads per sample. Analysis of the transcripts was performed using featureCounts (Liao et al. 2014) and differentially expressed genes (DEGs) were called with DESeq2 (Love et al. 2014) using R version 4.3.1 (Team 2023) and R Studio (Team 2022). Transcripts were identified as statistically significant with a Benjamini-Hochberg (BH) adjusted p value < 0.05 and absolute log2 fold change (log_2_FC) ≥1.

Principal component analysis was performed within DESeq2 using the function plotPCA on the 1,000 most variable genes. Further analyses were carried out with the normalized counts generated by DESeq2, which applies the median-of-ratios method to compute a size factor per sample, thereby accounting for sequencing depth and RNA composition. Normalized counts correspond to raw counts divided by these size factors. Heatmaps were generated using the R package pheatmap (Kolde 2025), based on the top 50 differentially expressed genes and normalized count values. Unsupervised hierarchical clustering was performed for rows. Counts were expressed as z-scores according to the equation (Eq. 1):

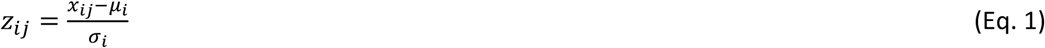

Where *x_ij_* is the normalized expression value of gene i in sample j, *μ_i_* is the mean expression of gene i across all samples, and *σ_i_* is the corresponding standard deviation.

Gene Ontology (GO) analysis was performed on DEGs using the R package clusterProfiler (Wu et al. 2021) with the mouse mm10 reference genome. Genes meeting standard significance cutoffs (adjusted p-value < 0.05 and |log_2_ fold change| ≥ 1) were included. Enrichment analysis was conducted separately for biological process (BP), molecular function (MF), and cellular component (CC) categories. GO terms with adjusted p-values < 0.05 were considered significantly enriched. Results were visualized using dotplot and cnetplot functions in clusterProfiler (Wu et al. 2021). Category–gene network plots (cnetplots) were generated to visualize the top five enriched GO terms and their associated genes. In these plots, nodes represent either GO terms or genes, and edges connect genes to the functional categories in which they are involved. This visualization highlights both the key biological processes enriched in the dataset and the overlap of genes contributing to multiple categories.

KEGG pathway enrichment was performed using the enrichKEGG function in clusterProfiler (Wu et al. 2021) with the same gene set. Pathways with adjusted p-values < 0.05 were considered significant.

### Data Availability

The RNA-sequencing data generated in this study have been deposited in the NCBI Sequence Read Archive (SRA) under BioProject accession no. PRJNA1530437. Custom scripts are available on GitHub at https://github.com/angelikaLahn/surface_curvature under the MIT License.

## Results

The experimental design allowed time-dependent as well as geometry-dependent comparison of gene expression in murine pre-osteoblastic MC3T3-E1 cells that were cultured on curved and flat surfaces, respectively, for up to 23 days (Fig. 1A). In total 2,124 DEGs were identified for CB and 1,166 for FT meeting the cutoff of absolute log_2_FC ≥ 1 and a BH p < 0.05. Of these transcripts, 619 are shared between the two culture conditions, whereas 1,505 are unique for CB and 546 for cells grown on FT surfaces (Fig. 1B, Tab. S1). Principal component analysis (PCA) based on the top 1,000 differentially expressed genes (DEGs) revealed distinct clustering of samples according to the surface condition and culture duration (Fig. 1C). Notably, samples from cells grown on flat surfaces for 23 days did not cluster well, likely due to over-confluence and senescence-associated cellular stress, which was also evident in the differential gene expression analysis. When comparing day 23 versus day 7, 28 senescence-related genes were identified in CB cultures and 21 in FT cultures, of which 15 genes were shared, indicating a common core of senescence genes activated over time (Tab. S2). In contrast, comparison between CB and FT surfaces at the same time points showed fewer shared genes: suggesting that the activation of senescence pathways is largely dependent on substrate geometry, with CB surfaces inducing a broader senescence response over the culture period. Overall, CB cultures showed a broader senescence response, with 16 upregulated and 5 downregulated senescence-related genes among 2,124 DEGs, compared with 6 upregulated and 5 downregulated genes among 1,165 DEGs in FT cultures.

### Time-dependent gene expression on curved surfaces

On curved surfaces a time-dependent expression of genes was observed showing up- and down-regulated genes with a log_2_ fold change of 1 and 2 (log_2_FC) and a statistical significance p < 0.05 (-log_10_(p)>1.3) comparing gene’s expression change of cells grown on capillary bridges at day 23 compared to day 7 (Fig. 2A, Tab. S1). Volcano plots showed that the top significantly upregulated genes were associated with signaling receptor binding and co-receptor activities, like *Serpine1* (serine/cysteine peptidase inhibitor clade E member1), *Ig@p2* (insulin-like growth factor binding protein 2), *Ramp1* (receptor (calcitonin) activity modifying protein 1), *Trf* (transferrin), *Sstr2* (somatostatin receptor 2), and *Itgbl1* (integrin beta-like 1). A hormone receptor, *Nr4a3* (nuclear receptor subfamily 4, group A, member 3), that binds to DNA and modulates gene expression (Ma et al. 2024) was also significantly upregulated after 23 days on curved surfaces. Conversely, the strongest downregulation was observed with *Ccn3* (cellular communication network factor 3) known to regulate osteoblast differentiation. Genes involved in cell proliferation (*Enpep* (glutamyl aminopeptidase)) and receptor signaling (*Plxna2* (plexin A2)) were also significantly downregulated (Fig. 2A).

**Figure 2.**
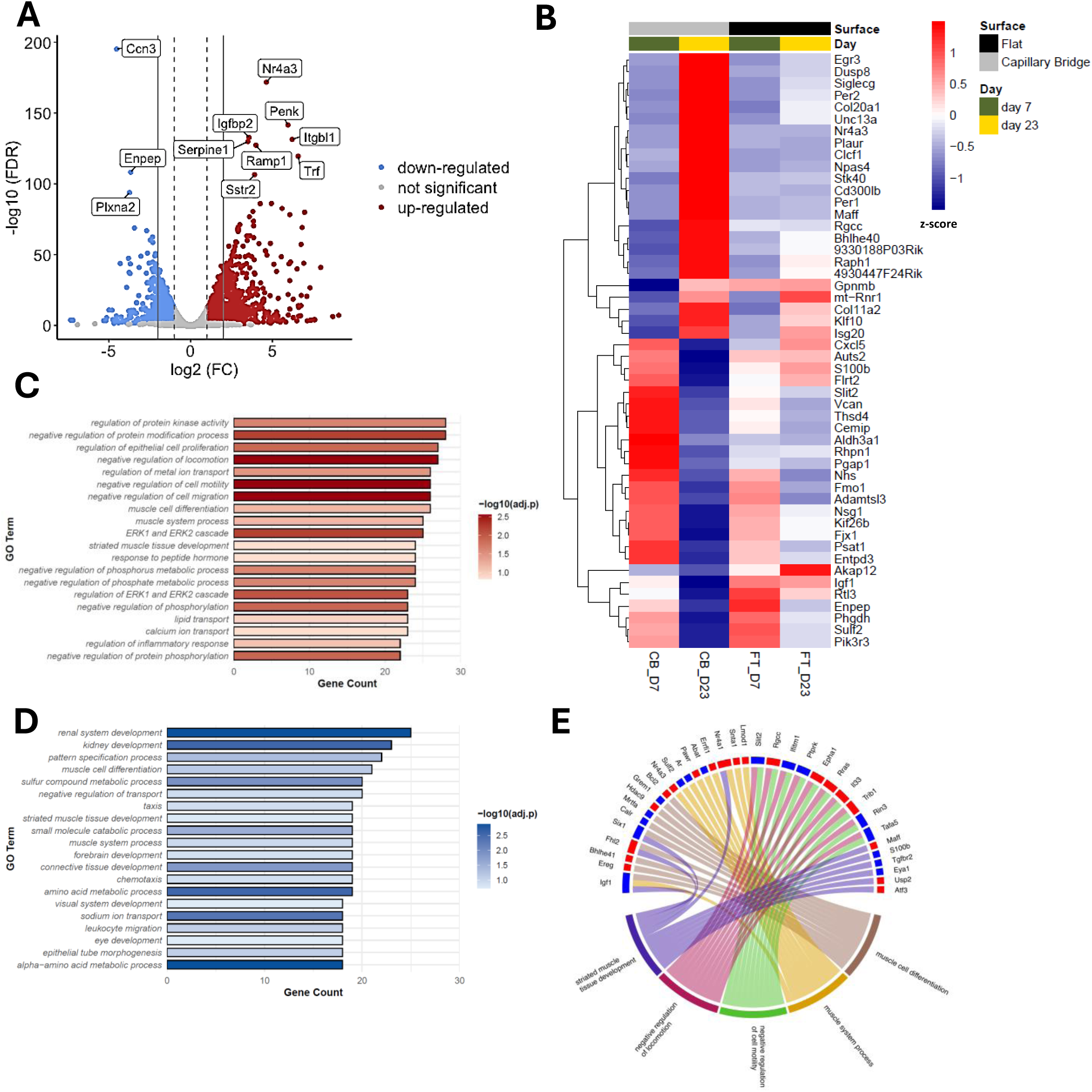
Time-dependent gene expression on curved surfaces. **A.** Volcano plot visualising the up- and down-regulated genes of cells grown on CB surfaces at day 23 compared to day 7. Dashed grey lines show cut-offs at log2 fold change ≥1 or ≤-1 corresponding to a doubling of values, whereas solid grey lines show a cut-off at log2 fold change ≥2 or ≤-2. The y-axis indicates the statistical significance as the log10 of the false discovery rate (FDR) corrected p-values. **B.** A heatmap shows the hierarchical clustering of the top 50 differentially expressed genes (DEGs) comparing 23-days-old against 7-days-old cells grown on curved surfaces. Z-score values of DEGs for cells cultured on flat (FT, black bar) and curved (CB, grey bar) surfaces for 7 (green bar) and 23 (yellow bar) days, respectively, are shown. Mean z-scores from replicates of the 4 conditions are given. **C.** The top 20 up-regulated biological processes in cells grown on curved surfaces after 23 days. Unique DEGs were sorted to Gene Ontology categories (GO terms) **D.** The top 20 down-regulated biological processes in cells grown on curved surfaces after 23 days. Unique DEGs were sorted to Gene Ontology categories (GO terms) **E.** Top 5 biological processes of all unique DEGs of cells grown on curved surfaces at day 23 compared to day 7. Individual DEGs of the same biological processes can be either up-(red) or down-regulated (blue).

Hierarchical clustering analysis showed that the top 50 unique DEGs comparing 23-days-old against 7-days-old cells grown on curved surfaces were shared with the top 50 DEGs of cells grown on FTs (Fig. 2B). However, cells on CBs showed an upregulation in time of transcriptional regulators, including transcription factors (*Egr3* (early growth response 3), *Npas4* (neuronal PAS domain protein 4), *Bhlhe40* (basic helix-loop-helix family, member 40), and *Klf10* (Kruppel-like transcription factor 10)) and genes involved in positive and negative regulation of transcription (*Per1* and *Per2* (period circadian clock 1 and 2), *Nr4a3* (nuclear receptor subfamily 4, group A, member 3), and *Maff* (v-maf musculoaponeurotic fibrosarcoma oncogene family protein F)). Another subset of DEGs showed elevated expression of genes associated with signaling-related processes, such as response to stimulus (*Dusp8* (dual specificity phosphatase 8), regulation of calcium-mediated signaling (*Siglecg* (sialic acid binding Ig-like lectin G)), and signaling receptor binding activities (*Unc13a* (unc-13 homolg A), *Plaur* (plasminogen activator, urokinase receptor), *Clcf1* (cardiotrophin-like cytokine factor 1), *Stk40* (serine/threonine kinase 40), *Cd300lb* (CD300 molecule like family member B), *Rgcc* (regulator of cell cycle), *Raph1* (Ras association (Ral GDS/AF-6 and pleckstrin homology domains 1), and *Gpnmb* (glycoprotein (transmembrane) nmb)). In addition, several extracellular matrix–associated genes, most notably *Col20a1* (collagen type XX alpha 1) and *Col11a2* (collagen type type XI alpha 2), were among the highest amounts of transcripts after 23 days.

On the other hand, a set of genes associated with signaling receptor binding (*Cxcl5* (C-X-C motif chemokine ligand 5), *S100b* (S100 protein, beta polypeptide), *Flrt2* (fibronectin leucine rich transmembrane protein 2), *Nsg1* (neuron specific gene family member 1), *Igf1* (insulin-like growth factor 1), and *Pik3r3* (phosphoinositide-3-kinase regulatory subunit3)), and genes involved in signaling (*Auts2* (autism susceptibility candidate 2), *Fjx1* (four jointed box 1)) exhibited pronounced downregulation on day 23 on curved surfaces (Fig. 2B). Also, genes involved in cell proliferation (*Enpep* (glutamyl aminopeptidase), *Phgdh* (3-phosphoglycerate dehydrogenase), and *Sulf2* (sulfatase 2) and *Kif26b* (kinesin family member 26 B), showing microtubule binding and motor activity, were strongly downregulated indicating a complex response in the time-dependent gene expression of cells.

This was confirmed by the expression of genes clustered to specific ontologies (GO terms, Figs. 2C and 2D) which showed highly significant upregulation of gene categories related to negative regulation of cell migration and motility, consistent with a late differentiation state characterized by reduced cellular movement and increased matrix anchoring. Concurrently, an elevated enrichment of ERK1/2 signaling and phosphorylation-related processes suggested ongoing fine-tuning of signaling pathways required for osteoblast maturation. A significant downregulation of gene expression profiles showed a subset of GO terms related to renal system and kidney development, as well as metabolic and transport processes, including amino acid metabolism and sodium ion transport (Fig. 2D). The GO terms enrichment of renal system and kidney development were not limited to renal tissue but implied that many genes annotated to these processes encode broadly expressed transporters, metabolic enzymes, and regulatory factors that are shared across multiple tissues and developmental stages.

The top five biological processes (GO terms) enriched in DEGs at day 23 revealed strong associations with muscle-related processes (striated muscle tissue development, muscle system process, and muscle cell differentiation) as well as negative regulation of cell motility and of cell locomotion (Fig. 2E). Individual genes contributing to each process revealed genes within the same process were up-(red) or down-regulated (blue) thus, indicating individual regulation patterns for single genes within each functional category. For completeness, a similar data analysis was performed for the time-dependent gene expression on flat surfaces (Fig. S1) but will not be considered furthermore in this study.

### Surface-dependent gene expression

Expression analysis between flat and curved surfaces at day 7 (Fig. 3A) revealed a pronounced early transcriptional response to surface curvature. Among the most significantly upregulated genes were early transcription factors *Egr1* (early growth response 1) and *Egr2* (early growth response 2), *Junb* (jun B proto-oncogene) as well as *Nr4a1* (nuclear receptor subfamily 4, group A, member 1), *Btg2* (BTG anti-proliferation factor 2) and *Cyp1b1* (cytochrome P450, family 1, subfamily b, polypeptide 1), *Crlf1* (cytokine receptor-like factor 1) and *Aldh1l2* (aldehyde dehydrogenase 1 family, L2). These genes are classically associated with rapid activation due to mechanical stimulation and stress-responsive signaling pathways as well as early differentiation processes. In contrast, genes associated with inflammatory signaling and extracellular matrix (ECM) remodeling, including *Il33* (interleukin 33), *Gpnmb* (glycoprotein (transmembrane) nmb) and *F13a1* (coagulation factor XIII, A1 subunit) were strongly downregulated on curved surfaces suggesting a curvature-induced activation of mechanosensitive transcriptional programs while suppressing inflammatory and remodeling pathways at this initial stage.

**Figure 3.**
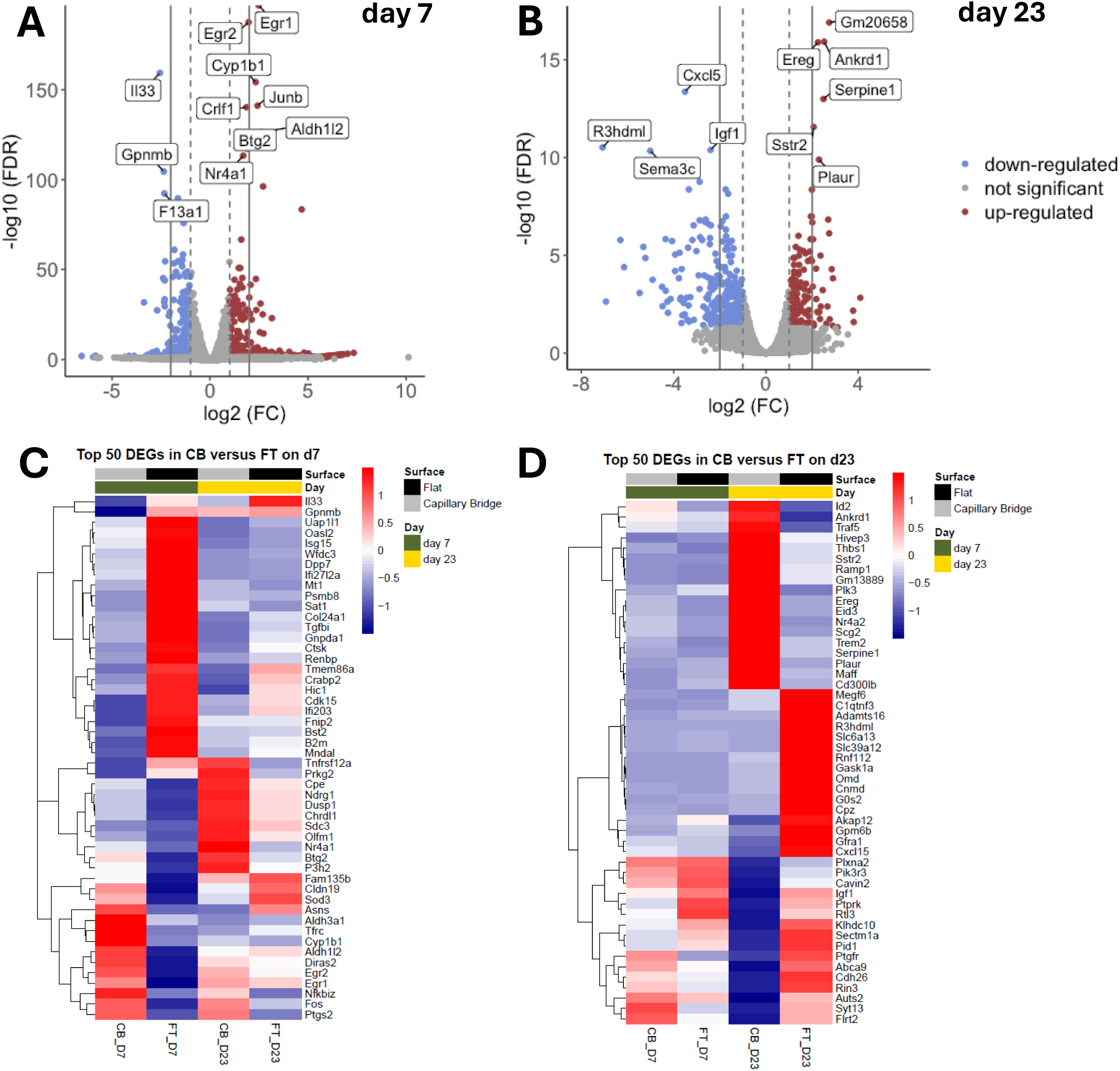
Surface-dependent gene expression. **A.** Volcano plot shows differential gene expression in curved surfaces (CB) compared to flat surfaces (FT) for day 7 and **B.** for day 23. Dashed grey lines show cut-offs at log2 fold change ≥1 or ≤-1 corresponding to a doubling of values, whereas solid grey lines show a cut-off at log2 fold change ≥2 or ≤-2. The y-axis indicates the statistical significance as the log10 of the false discovery rate (FDR) corrected p-values. **C.** Heatmap of the top 50 uniquely DEGs comparing cells grown on curved surfaces against cells grown on flat surfaces after 7 days. Z-score values of DEGs for cells cultured on flat (FT, black bar) and curved (CB, grey bar) surfaces for 7 (green bar) and 23 (yellow bar) days, respectively, are shown. Mean z-scores from replicates of the 4 conditions are given. **D.** Heatmap of the top 50 uniquely DEGs comparing cells grown on curved surfaces against cells grown on flat surfaces after 23 days. Z-score values of DEGs for cells cultured on flat (FT, black bar) and curved (CB, grey bar) surfaces for 7 (green bar) and 23 (yellow bar) days, respectively, are shown. Mean z-scores from replicates of the 4 conditions are given.

Later on, at day 23, the transcriptional pattern on curved surfaces shifted toward genes associated with ECM organization and growth factor signaling (Fig. 3B). The most significantly upregulated genes were *Ankrd1* (ankyrin repeat domain 1), *Ereg* (epiregulin), *Sstr2* (somatostatin receptor 2), *Serpine1*(serine (or cysteine) peptidase inhibitor, clade E, member 1) and *Plaur* (plasminogen activator, urokinase receptor). These genes are involved in ECM remodeling, mechanotransduction, EGFR-mediated signaling and proteolytic regulation. In contrast, *Cxcl5* (C-X-C motif chemokine ligand 5), *Igf1* (insulin-like growth factor 1), and *Sema3c* (semaphorin 3 C), were markedly downregulated, suggesting modulation of chemokine signaling and growth factor pathways.

The expression data of the top 50 unique DEGs revealed a distinct early transcriptional response to curvature (Fig. 3C). Curved surfaces induced upregulation of immediate early transcription factors and stress-responsive genes, including again *Egr1* and *Erg2* (early growth response 1 and 2), *Fos* (Fos proto-oncogene, AP-1 transcription factor subunit), *Nffibiz* (nuclear factor of kappa light polypeptide gene enhancer in B cells inhibitor, zeta), and *Ptgs2* (prostaglandin-endoperoxide synthase 2), together with metabolic regulators such as *Asns* (asparagine synthetase), *Aldh1l2* and *Ald3a1* (aldehyde dehydrogenase 1 family L2 and family 3, subfamily A1), and *Cyp1b1* (cytochrome P450, family 1, subfamily b, polypeptide 1). In contrast, pronounced downregulation was observed for genes associated with inflammatory signaling and immune-related pathways, including *Il33* (interleukin 33), *Gpnmb* (glycoprotein (transmembrane) nmb), also known as osteoactivin, as well as *B2m* (beta-2 microglobulin), *Bst2* (bone marrow stromal cell antigen 2), *Tnfrsf12a* (tumor necrosis factor receptor superfamily, member 12a), and *Ifi203* (interferon activated gene 203). Several additional genes involved in interferon responses, extracellular matrix regulation, and differentiation markers showed moderate downregulation, while *Cldn19* (claudin 19) and *Sod3* (superoxide dismutase 3) were slightly upregulated. Together, these data indicate a curvature-dependent emphasis on transcriptional and stress-responsive gene expression on day 7.

During the late culture phase on curved surfaces (day 23, Fig. 3D) a subset of genes associated with signal transduction and receptor-mediated pathways exhibited significant upregulation, including *Sstr2* (somatostatin receptor binding), *Ramp1* (receptor (calcitonin) activity modifying protein1), *Ereg* (epiregulin), and *Traf5* (TNF receptor-associated factor 5) as well as genes involved in ECM remodeling (*Serpine 1* (serine/cysteine peptidase inhibitor, clade E, member 1) and *Plaur* (plasminogen activator, urokinase receptor-related)) and interactions between cell and matrix, like *Thbs1* (thrombospondin 1). Furthermore, genes for mechanosensitive transcription regulation, especially *Ankrd1* (ankyrin repeat domain 1), or *Hivep3* (human immunodeficiency virus type I enhancer binding protein 3), *Eid3* (EP 300 interacting inhibitor of differentiation 3), *Maff* (v-maf musculoaponeurotic fibrosarcoma oncogene family protein F), and *Id2* (inhibitor of DNA binding 2), and a group of genes involved in immune-associated signaling, including *Trem2* (triggering receptor expressed on myeloid cells 2), and *Cd300lb* (CD 300 molecule like family member B) were upregulated. On the other hand, downregulated genes were predominantly detected in categories of intracellular signaling pathways, like *Pik3r3* (phosphoinositide-3-kinase regulatory subunit 3), *Igf1* (insulin-like growth factor), *Ptgfr* (prostaglandin F receptor), *Ptprk* (protein tyrosine phosphatase receptor type K), *Rin3* (Ras and Rab interactor 3), and *Pid1* (phosphotyrosine interaction domain containing 1), and cell adhesion, like *Plxna2* (plexin A2), *Cdh26* (cadherin-like 26), and *Flrt2* (fibronectin leucine-rich transmembrane protein 2), or membrane dynamics, like *Cavin2* (caveolae associated 2), *Syt13* (synaptotagmin XIII), and *Abca9* (ATP-binding cassette, subfamily A member 9) with additional contributions from transcriptional regulation, like *Auts2* (autism susceptibility candidate 2), *Rtl3* (retrotransposon Gag like 3), oxidative stress (*Klhdc10* (kelch domain containing 10)), and immune-associated processes (*Sectm1a* (secreted and transmembrane 1A)). Additionally, a moderate downregulation showed a group of genes involved in ECM organization, such as *Adamts16* (ADAM metallopeptidase with thrombospondin type 1 motif 16), *Cpz* (carboxypeptidase Z), *Megf6* (multiple EGF-like-domains 6) *C1qtnf3* (C1q and tumor necrosis factor related protein 3) and bone/cartilage remodeling, like *Cnmd* (chondromodulin) and *Omd* (osteomodulin).

### Curved surfaces regulate major biological processes in pre-osteoblasts

To arrange these complex expression patterns of individual genes into biological processes, a GO term enrichment analysis was performed. At day 7, up-regulated GO terms in cells grown on curved compared to flat surfaces were predominantly associated with early cellular responses, including response to metal ions, response to mechanical stimulus, cell–cell adhesion via plasma membrane adhesion molecules, and processes related to synapse assembly and neuronal-like signaling (Fig. 4A). Cells on day 23 (Fig. 4B) showed an increase in up-regulated genes belonging to GO terms of tissue maturation and differentiation, striated muscle tissue development, muscle organ development, and skeletal muscle tissue development. The enrichment in muscle-related GO terms likely reflects cytoskeletal remodeling, contractility and mechanotransduction processes that are commonly linked to “muscle development genes” rather than cell lineage-specific differentiation. In contrast, reduction of gene expression was observed on day 7 in processes associated with antiviral and immune-related pathways, including defense response to virus, regulation of viral process, and regulation of viral genome replication (Fig. 4C). The down-regulation of antiviral and interferon-associated pathways on day 7 may reflect reduced stress- or inflammation-related signaling during early adaptation to curved substrates. At a later growth phase (day 23) cells exhibited down-regulation of processes related to taxis and chemotaxis, extracellular matrix (ECM) organization, extracellular structure organization, developmental growth involved in morphogenesis, developmental cell growth, and renal/kidney development-associated pathways (Fig. 4D). This down-regulation at day 23 may indicate that migration decreases and cells turned into a sessile steady-state, thus reflecting a transition between physiological states from day 7 to day 23.

**Figure 4.**
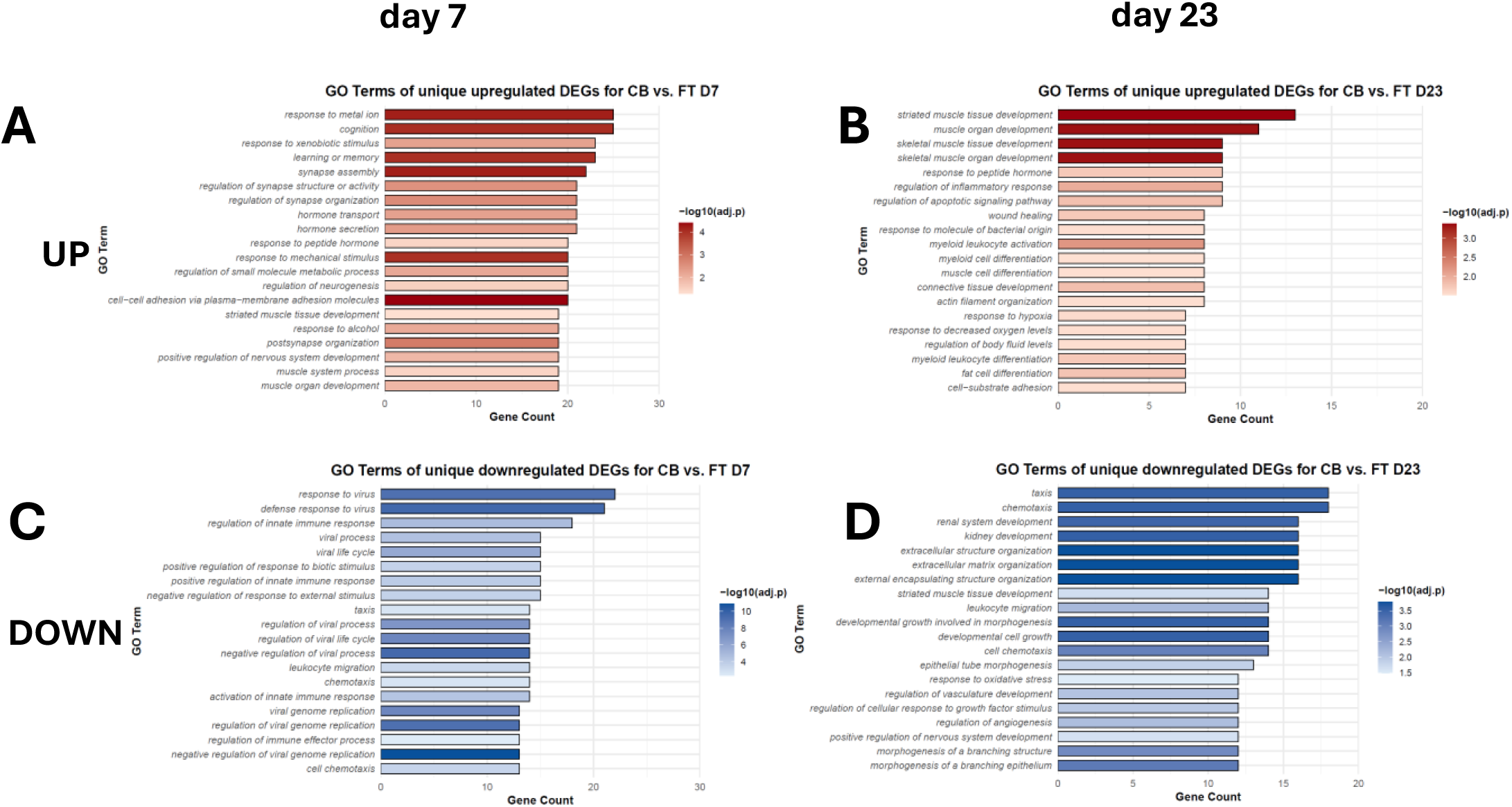
Major biological processes in curved surface-grown cells. **A.** The top 20 up-regulated biological processes in cells grown on curved surfaces for 7 days compared to cells grown on flat surfaces of same culture age. Unique DEGs were sorted to Gene Ontology categories (GO terms). **B.** Top 20 up-regulated biological processes in cells grown on curved surfaces for 23 days compared to cells grown on flat surfaces of same culture age. Unique DEGs were sorted to Gene Ontology categories (GO terms). **C.** Top 20 down-regulated biological processes in cells grown on curved surfaces for 7 days compared to cells grown on flat surfaces of same culture age. Unique DEGs were sorted to Gene Ontology categories (GO terms). **D.** Top 20 down-regulated biological processes in cells grown on curved surfaces for 23 days compared to cells grown on flat surfaces of same culture age. Unique DEGs were sorted to Gene Ontology categories (GO terms).

### Curvature modulates signaling pathways of cellular adaptation, proliferation, and differentiation

To further obtain insight into the molecular mechanisms by which cells respond to curvature, the expression of specific genes known to be involved in osteoblast differentiation were analyzed (Salhotra et al. 2020, Zhu et al. 2024). As shown in Fig. 5A cells cultured on curved relative to flat surfaces revealed almost no changes (Δlog_2_FC ≤ 1) of osteogenic transcription factors, like *Sox9* (SYR-box transcription factor 9), *Runx1* and *2* (Runt-related transcription factor 1 and 2), *Cb@* (core-binding factor subunit beta), *Osx/Sp7* (Osterix), *Atf4* (activating transcription factor 4), and *Yap1* (Yes-associated protein 1, whereas bone morphogenic signaling molecules exhibited pronounced temporal changes: *Bmp2* (bone morphogenetic protein 2) expression increased from day 7 to day 23 by a Δlog_2_ FC of 4 and *Bmp1b* (bone morphogenetic protein receptor type 1B) showed a decrease of log_2_FC of 3 (Fig.5B). In addition, the Notch target genes *Hey1* and *2* (hairy/enhancer-of-split related with YRPW motif 1 and 2), and *Hes3* (hes family bHLH transcription factor 3) increased from slight negative values of log_2_FC ≈ −1 at day 7 to positive values of log_2_FC ≈ 0.5–2 on day 23 (Fig. 5C). In Wnt signalling pathways the expression of several genes, e.g. *Wnt8b* (wingless-type MMTV integration site family, member 8B) and *Fzd3* (Frizzled receptor 3) decreased, while *Wnt9b* (wingless-type MMTV integration site family, member 9B) increased from clearly down-regulated to up-regulated, from log_2_FC = −3 to log_2_FC = +3 (Fig. 5D). Among Fibroblast Growth Factor family members, expression of *Fgf1*, *Fgf14*, *Fgf21*, and *Fgf23* decreased between day 7 and day 23, whereas *Fgf13* and *Fgf20* showed increased expression, ranging from approximately −2 to +3 (Fig. 5E). These data indicate that substrate curvature primarily affected growth factors and signaling pathway-associated genes rather than core osteogenic transcription factors.

**Figure 5.**
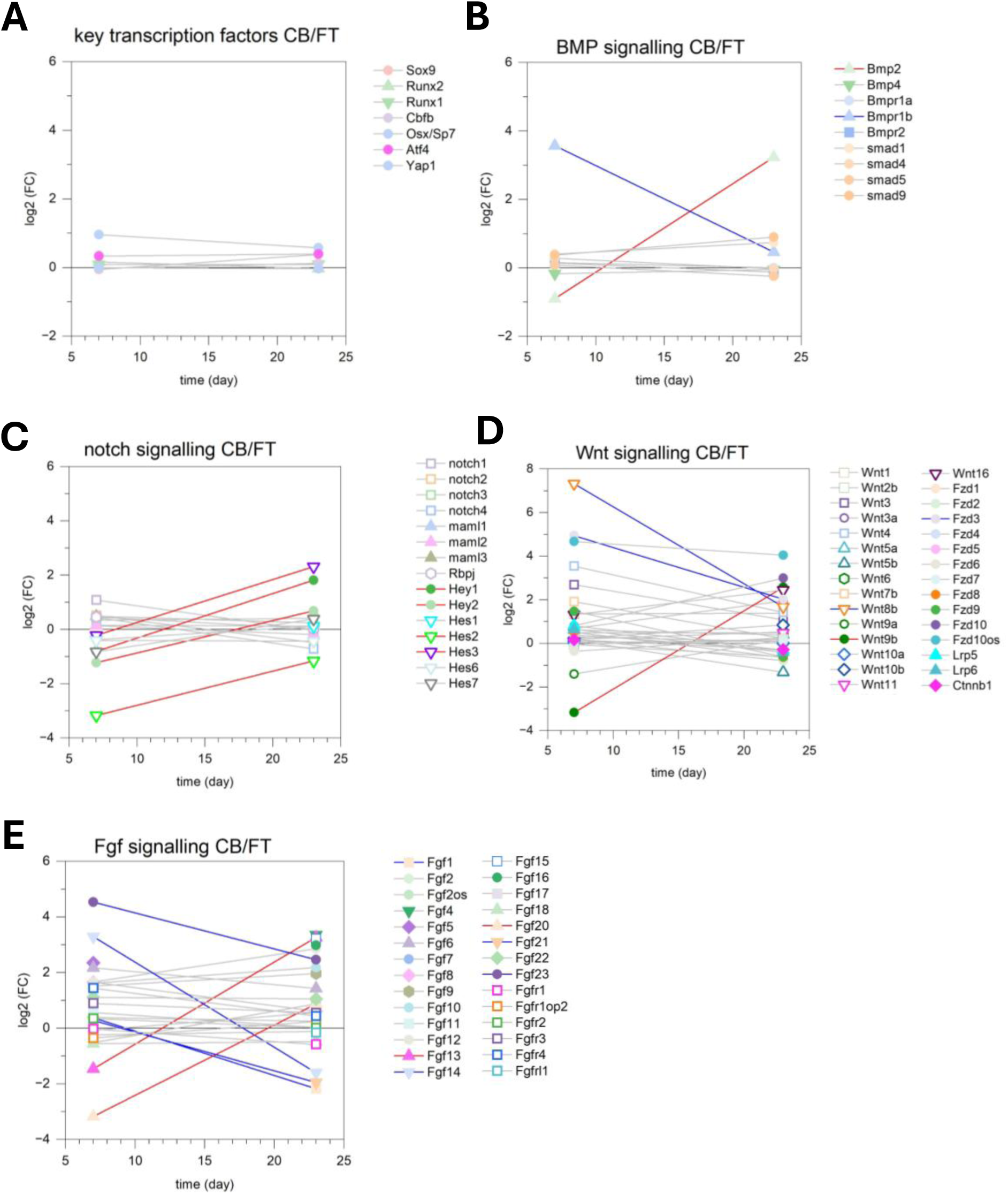
Osteoblast differentiation. Changes in key transcription factors and genes of major signalling pathways (according to Salhotra et al., 2020 and Siyu Zhu et al., 2024) from cells grown on curved surfaces compared to flat surfaces are plotted against the cultivation time. **A.** Transcription factors, **B.** Bone morphogenetic protein (BMP) signalling, **C.** Notch signalling, **D.** Wnt signalling and, **E.** Fibroblast growth factor (FGF) signalling and Larger changes between day 7 and day 23 with log2 ≥ 2 and ≤ −2 are marked by red (up) and blue (down) lines, respectively. Data from supplemental Table S1.

### Curved substrates promote dynamic remodeling of the extracellular matrix

A KEGG pathway enrichment analysis for “ECM-receptor interaction” (KEGG ID: mmu04512) showed unique DEGs that were significantly differentially expressed on curved surfaces on day 7 (Fig. 6A) and day 23 (Fig. 6B) mainly including collagens, laminins, and integrins. Among these, the collagen *Col26a1* exhibited a strong change of expression over 6 magnitudes of log_2_, from upregulation at day 7 (log_2_ FC ≈ 4.2) followed by downregulation at day 23 (log_2_ FC ≈ −2.2). A similar but moderate change was observed with *Col9a2*: upregulated at day 7 (log_2_ FC ≈ 1.2) and downregulated at day 23 (log_2_ FC ≈ −1.0). In contrast, *Col9a3* showed weak downregulation at day 7 but increased to upregulation on day 23, indicating a rebuilding of the ECM during prolonged culture on curved substrates (Fig. 6C). Most laminin genes (*Lam a1 - Lam a5*, *Lam b1, Lam b3* and *Lam c1*) displayed only slight, decreasing changes in expression (Fig. 6D), whereas Lam b2and *Lam c2* showed a slight increase on curved compared to flat surfaces (Fig. 6D). Strong temporal changes were detected for ECM-associated proteins (Fig. 6E). Vitronectin (*Vtn*) and thrombospondin 4 (*Thbs4*) shifted from strongly downregulated at day 7 to upregulated at day 23 with a Δlog_2_FC of 6 and 4.5, respectively. Conversely, *Spp2* (secreted phosphoprotein 2) changed from moderate upregulation at day 7 (log_2_ FC ≈ 2) to downregulation at day 23 (log_2_ FC ≈ −2). Integrin genes likewise display dynamic regulation (Fig. 6F). *ItgaD* and *Itga8* changed from upregulation at day 7 (log_2_ FC ≈ 4) to less upregulated (Δlog_2_FC ≈ −2) and even downregulation (Δlog_2_FC ≈ −2.3), respectively, whereas the expression of other integrin genes like *ItgaM, Itga4* and ItgaX increased on curved surface from day 7 to day 23. Altogether, these data indicated that substrate curvature induced selective and time-dependent remodeling of the extracellular matrix by precisely changing the components and their associated adhesion receptors.

**Figure 6.**
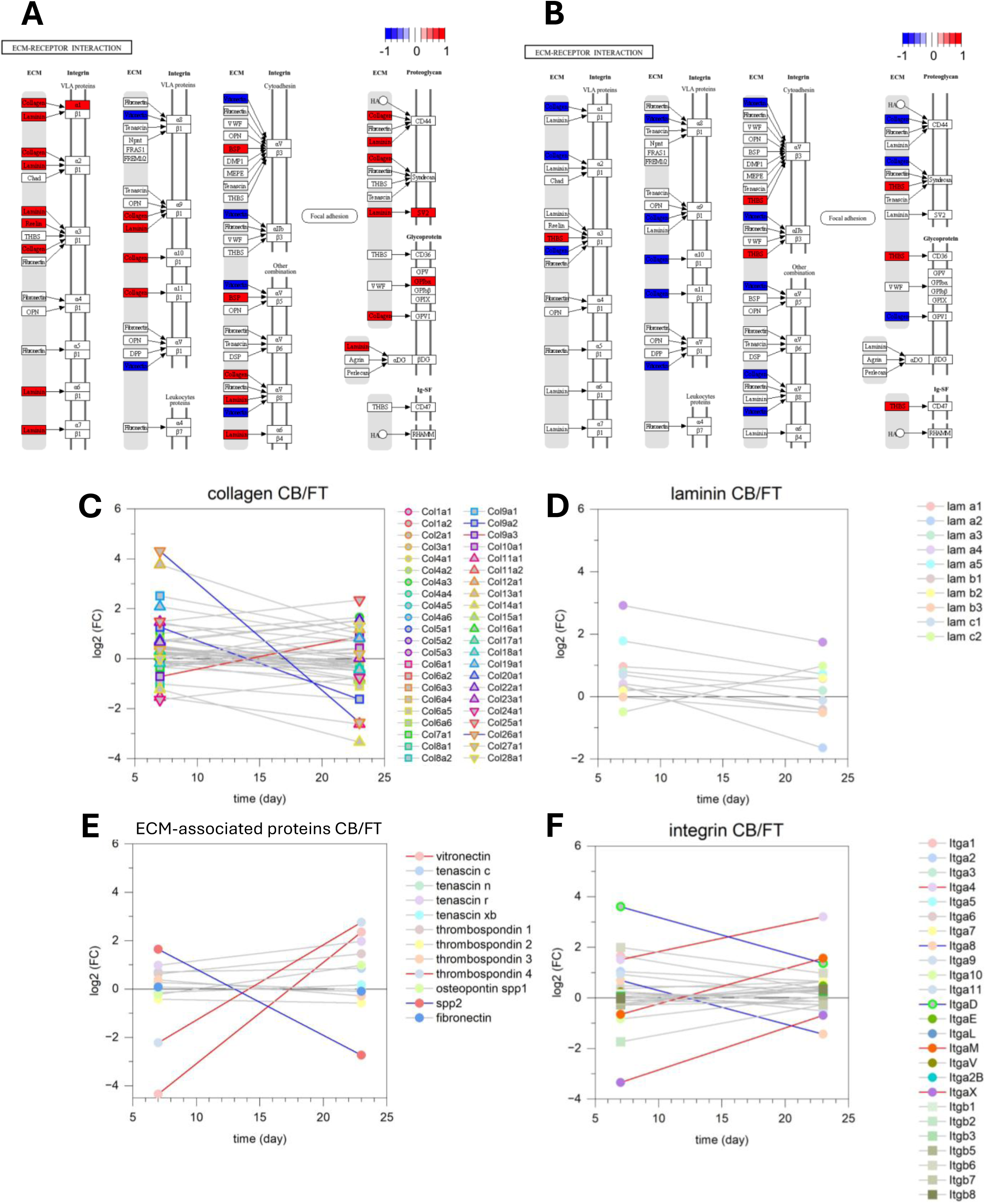
Surface-dependent ECM-cell receptor interaction. KEGG pathway (mmu04512) enrichment analysis of significant (Benjamini-Hochberg adjusted p < 0.05), unique DEGs from cells grown on curved surfaces compared to flat surfaces was performed for **A.** day 7 and **B.** day 23. Red coloured genes are up-regulated and blue coloured are down-regulated. **C**. collagens, **D**. laminins, **E.** ECM-associated proteins, and **F**. integrins. Larger changes between day 7 and day 23 are marked by blue (down) and red (up) lines. Data from supplemental Table S1.

Inside the cell, the cytoskeleton may transduce the signals of the ECM from the curved environment. Therefore, cytoskeletal adaptations, differentially expressed genes associated with actin organization and cytoskeletal regulation were analyzed (Fig. 7). Within the group of “actin filaments”, *Acta1* (actin, alpha 1) exhibited a marked increase in expression from a log_2_ fold change of approximately 0.2 at day 7 to approximately 3.0 at day 23. In contrast, *Acte1* (actin, epsilon 1) expression decreased from approximately 4.3 at day 7 to 1.8 at day 23. Other actin isoforms, including *Acta2* (actin, alpha 2), *Actb* (actin, beta), *Actbl2* (actin, beta-like 2), *Actb-ps1* (actin, beta, pseudogene 1), *Actc1* (actin, alpha, cardiac muscle 1), *Actg1* (actin, gamma, cytoplasmic 1), *Actg2* (actin, gamma 2), and *Actg-ps1* (actin, gamma, pseudogene 1), showed only minor changes over time (Fig. 7A). In the group of actin-binding proteins, the most strongly regulated genes, such as *Was* (Wiskott-Aldrich syndrome), *Wasf3* (WASP family, member 3), and *Fhod3* (formin homology 2 domain containing 3) exhibited decreased expression from day 7 to day 23, whereas *Coro6* (coronin 6) increased from slight downregulation (log_2_ FC ≈ −1.1) to moderate upregulation (log_2_ FC ≈ 0.4, Fig. 7B). Among genes associated with stress fibers, *Myh2* (myosin, heavy polypeptide 2) showed a pronounced increase in expression from approximately 2.0 to 4.9, while *Actn3* (actinin alpha 3) decreased from approximately 0.8 to −0.5 (Fig. 7C). Genes encoding for focal adhesion proteins, vinculin, filamin A, palladin, transgelin, and related proteins, displayed only modest expression changes (Fig. 7D). Similarly, genes associated with “intermediate filaments”, including among others *Lmna* (lamin A), *Lmnb1* (lamin B1), *Lmnb2* (lamin B2), desmin, vimentin, and nestin remained largely unchanged throughout the culture period (Fig. 7E). These findings indicate that substrate curvature predominantly affects genes involved in actin filament remodeling and contractile function, whereas focal adhesion and intermediate filament components exhibit comparatively stable expression.

**Figure 7.**
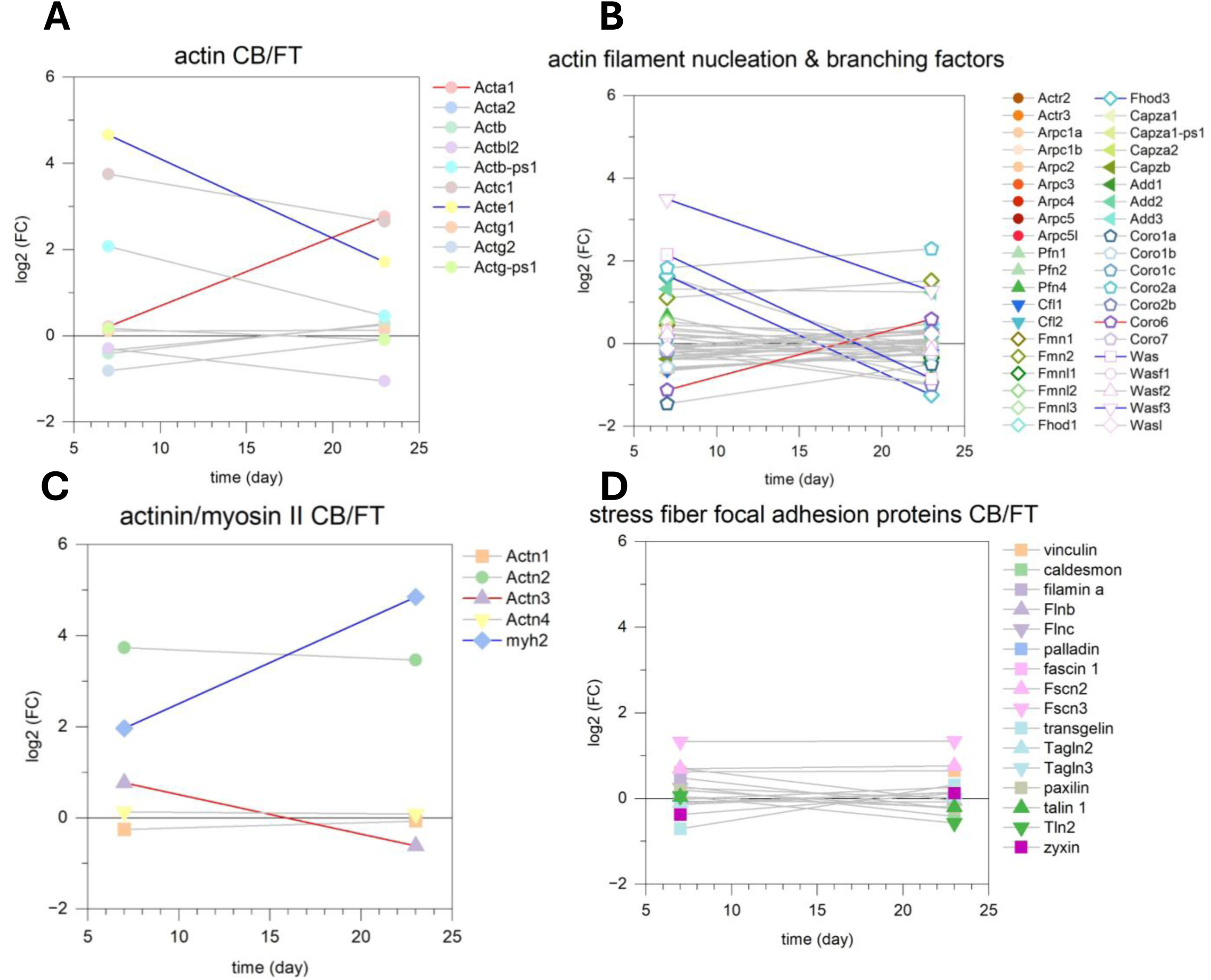
Surface-dependent expression of actin cytoskeleton and stress fibers. Unique DEGs from cells grown on curved surfaces were compared to cells grown on flat surfaces. Larger changes between day 7 and day 23 are marked by blue (down) and red (up) lines. Data from supplemental Table S1. **A.** Actin filaments **B.** Actin binding proteins, nucleation and branching factors **C.** Stress fibers: myosin II and actinin **D.** focal adhesion proteins

Analysis of known mechanosensitive cell components like PIEZO ion channels revealed only modest changes in expression of *Piezo1* and *Piezo2* (piezo-type mechanosensitive ion channel component 1 and 2) between curved and flat substrates (Fig. 8A). *Piezo1* expression remained largely unchanged throughout the culture period. *Piezo2* was consistently upregulated on curved surfaces but decreased from a log_2_ fold change of approximately 3.0 at day 7 to 2.0 at day 23.

**Figure 8.**
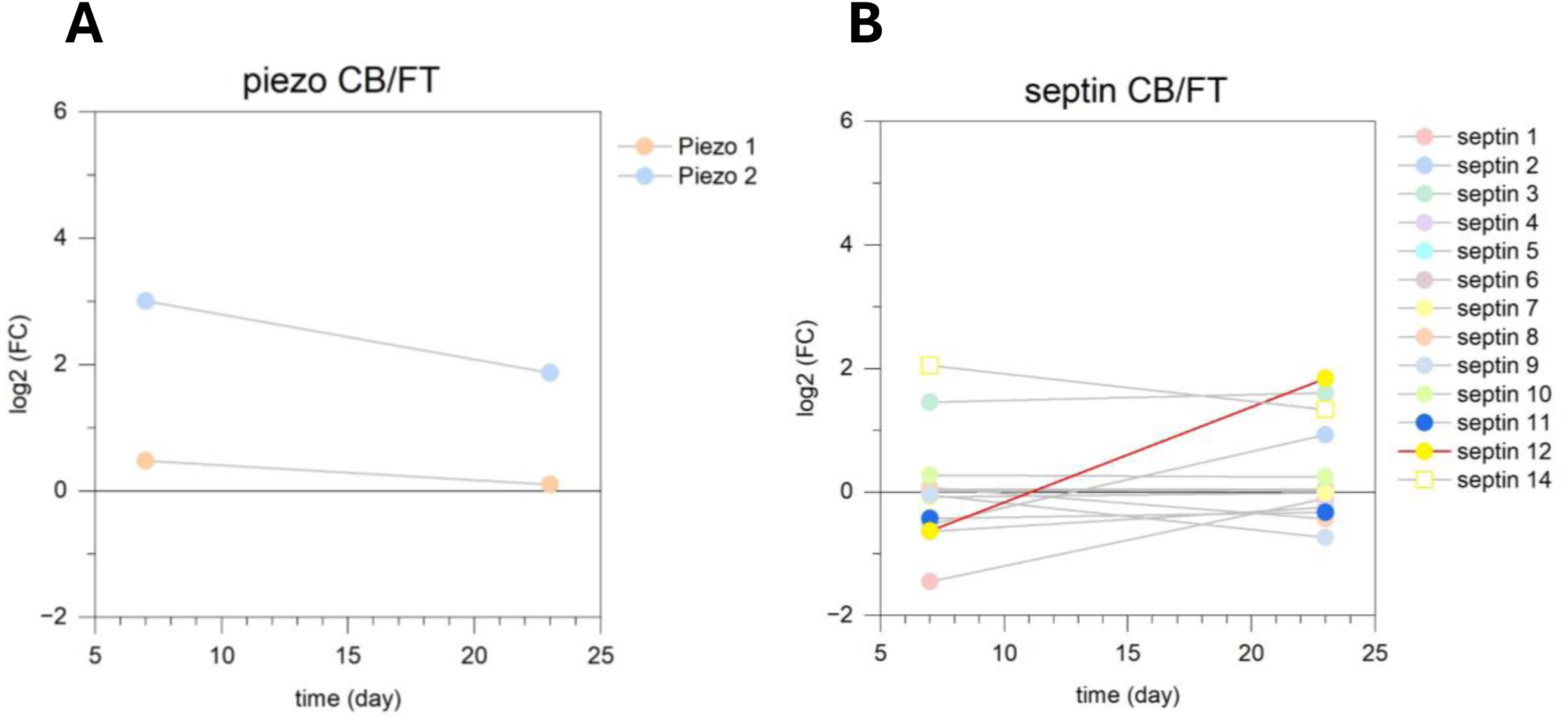
Surface-dependent expression of mechano-signaling components. Unique DEGs from cells grown on curved surfaces were compared to cells grown on flat surfaces. Larger changes between day 7 and day 23 are marked by blue (down) and red (up) lines. Data from supplemental Table S1. **A.** Mechanically activated ion channels Piezo 1 and 2 **B.** Septins

Expression analysis of the septin family, another mechanosensitive component, revealed only little transcriptional changes (Fig. 8B). Septin1–Septin11 and Septin14 exhibited only minor differences in expression between curved and flat substrates at both time points. In contrast, Septin12 showed a progressive increase in expression from slight downregulation on day 7 (log_2_ FC ≈ −0.6) to moderate upregulation on day 23 (log_2_ FC ≈ 2.0).

## Discussion

The present study demonstrates that substrate curvatures, with characteristic lengths much larger than that of a single cell, profoundly influence the transcriptional landscape of pre-osteoblastic MC3T3-E1 cells in a time-and surface-dependent manner. Principal component analysis (Fig. 1C) showed clear segregation according to both culture duration and substrate geometry, indicating that curvature affects not only the initial cellular adaptation but also long-term differentiation-associated transcriptional programs. Rather than directly altering the expression of the canonical osteogenic transcription factors *Runx2*, *Cb@*, *Osx*/*Sp7*, and *Atf4*, curvature primarily modulated extracellular matrix organization, cell–matrix adhesion, cytoskeletal remodeling, and interconnected BMP-, WNT-, FGF-, and Notch-dependent signaling pathways in a time-dependent manner. While prolonged culture induced a common set of senescence-related genes irrespective of the substrate, CB surfaces elicited a broader and more pronounced senescence-associated transcriptional signature than FT surfaces. The limited overlap in senescence-related genes between CB and FT cultures at identical time points further suggests that substrate curvature, rather than culture duration alone, is a major determinant of the transcriptional response.

### Time-dependent gene expression on curved surfaces

Comparison of day 23 with day 7 cultures on curved substrates revealed a marked shift from early adaptive responses toward signaling and differentiation-associated processes (Fig. 2). Increased expression of genes involved in receptor binding and signal modulation, including *Serpine1, Ig@p2, Ramp1, Trf, Sstr2,* and *Itgbl1*, together with the immediate-early transcription factor *Nr4a3*, indicates sustained responsiveness to extracellular and mechanical stimuli (McLatchie et al. 1998, Storch et al. 2001, Lahlou et al. 2004, Song et al. 2018, Mizukami et al. 2024, Meng et al. 2025). The concurrent strong upregulation of the immediate-early nuclear receptor *Nr4a3* suggests metabolic adaptation and sustained transcriptional responsiveness to environmental and mechanical cues (Boulet et al. 2022). In contrast, reduced expression of genes associated with proliferation and osteoblast development, including *Ccn3, Enpep,* and *Plxna2*, suggests progression toward a more differentiated cellular phenotype (Nogi et al. 2010, Holmes et al. 2017, Babey et al. 2024).

The heatmap of the 50 most differentially expressed genes (DEGs; Fig. 2B) further supports this transition by demonstrating coordinated upregulation of transcriptional regulators (*Egr3, Per1, Per2, Nr4a3, Npas4, Maff, Bhlhe40,* and *Klf10*) (Ooe et al. 2004, Kumbrink et al. 2010, Seuter et al. 2013, Tao et al. 2015, Katsuoka and Yamamoto 2016, Boulet et al. 2022, Luo et al. 2023) together with genes involved in receptor signalling, like *Plaur*, *Clcf1*, *Cd300lb*, *Raph1*, and *Gpnmb* (Furlan et al. 2007, Nahlé et al. 2019, Tsou and Sawalha 2020, Brazzo et al. 2021) and calcium-dependent pathways (*Siglecg,* (Hoffmann et al. 2007)) suggesting that curved surfaces may promote enhanced signal responsiveness coupled to transcriptional reprogramming. Upregulation of *Raph1* (lamellipodin) and *Gpnmb* (osteoactivin), both linked to cytoskeletal organization and cell–matrix interactions, like actin polymerization and integrin binding activity, respectively (Moussa et al. 2014, Bodo et al. 2017), together with increased expression of *Col20a1* and *Col11a2*, suggests enhanced extracellular matrix remodeling and structural maturation during prolonged culture. Conversely, coordinated downregulation of growth factor- and chemokine-associated genes, including *Igf1, Cxcl5, Pik3r3, Flrt2*, *Nsg1*, and *Pik3r3* (Steiner et al. 2005, Piazza et al. 2013, Hadem and Sharma 2017, Zhao et al. 2017, Shirakawa et al. 2019, Chen et al. 2020, Costa et al. 2021, Zhu and Chin-Sang 2024) together with genes involved in membrane remodeling and extracellular matrix organization (*Pgap1* and *Adamtsl3*) (Hall et al. 2003, Zoltewicz et al. 2009), indicates attenuation of proliferative signaling and stabilization of the cellular microenvironment. Notably, several genes associated with cell proliferation and metabolic activity, such as *Enpep*, *Phgdh*, and *Sulf2*, also exhibited reduced expression. The downregulation of *Kif26b*, a kinesin family member involved in microtubule binding and cytoskeletal organization during differentiation as well as in bone formation via modulating Wnt/ß-catenin signaling (Pickering et al. 2022, Yan et al. 2022), suggests additional alterations in cytoskeletal organization and intracellular signaling processes.

Gene ontology analysis further indicates that prolonged culture on curved substrates suppresses migratory programs while reinforcing pathways associated with protein phosphorylation and ERK1/2 signaling (Fig. 2C). Since ERK signaling promotes late osteoblast differentiation and matrix maturation (Kim et al. 2022), these findings suggest that curvature favors the acquisition of a mature, matrix-producing phenotype rather than continued proliferation or migration. Consistent with this interpretation, repression of amino acid metabolism and ion transport pathways (Fig. 2D) reflects the metabolic shift accompanying osteoblast maturation. Although genes within several functional categories were regulated in both directions (Fig. 2E), this pattern likely reflects selective remodeling of contractility- and motility-associated networks rather than generalized pathway activation, highlighting the finely tuned transcriptional adaptation induced by substrate curvature.

### Surface-dependent gene expression on curved surfaces

The substantially larger number of differentially expressed genes on curved substrates (Fig. 3A) demonstrates that substrate curvature exerts a sustained and distinct effect on the osteoblast transcriptional program. The temporal expression profiles indicate a transition from an early mechanosensitive response toward long-term structural adaptation.

At day 7, curvature predominantly induced immediate-early transcription factors, including *Egr1, Egr2, Junb,* and *Nr4a1*, which are well-established mediators of mechanotransduction and rapidly respond to altered cell geometry and cytoskeletal tension (Morawietz et al. 1999, Kumbrink et al. 2010, Bougault et al. 2012, Gaut et al. 2016). Simultaneous downregulation of *Il33,* an inflammatory mediator involved in bone homeostasis (Okragly et al. 2016) as well as *Gpnmb* with its integrin binding activity and involved in matrix remodeling (Moussa et al. 2014) suggests attenuation of inflammatory signaling and early extracellular matrix remodeling, consistent with transcriptional priming rather than immediate osteogenic matrix production. The strong upregulation of *Ankrd1*, a mechanosensitive gene, strongly induced by cytoskeletal tension and mechanical stress and a strong indicator of curvature-induced mechanical signaling (Samaras et al. 2015, Lopez et al. 2024), highlights sustained mechanotransduction under curved conditions. The corresponding heatmap (Fig. 3D) further demonstrates coordinated activation of mechanosensitive transcription factors (*Egr1, Egr2,* and *Fos*), inflammatory regulators (*Nffibiz* and *Ptgs2*), and metabolic adaptation pathways, while interferon- and immune-associated genes were broadly suppressed (Feng et al. 2023, Chen et al. 2025). Together, these findings indicate that substrate curvature initially triggers a rapid adaptive program that integrates mechanosensing with metabolic reprogramming while limiting inflammatory and stress-associated responses.

By day 23, the transcriptional profile shifted from early signaling toward extracellular matrix remodeling and structural adaptation (Fig. 3C). Upregulation of the mechanosensitive regulator *Ankrd1* provides strong evidence for sustained mechanical stimulation under curved conditions (Samaras et al. 2015, Lopez et al. 2024). Concurrent induction of *Serpine1, Plaur,* and *Ereg* further indicates activation of pathways involved in extracellular matrix turnover, proteolysis, and growth factor signaling (Komurasaki et al. 1997, Furlan et al. 2007, Mizukami et al. 2024). In addition, with the reduced expression of *Igf1, Cxcl5,* and *Sema3c* attenuation of proliferative, chemotactic, and morphogenetic signaling as cells progress toward a more mature and structurally stabilized phenotype is suggested.

The heatmap of day 23 cultures (Fig. 3E) supports this interpretation by revealing coordinated downregulation of genes involved in growth factor signaling (e.g. *Igf1*, *Pik3r3*, *Ptprk*), cell adhesion and membrane-associated components (e.g. *Plxna2*, *Cavin2*, *Flrt2*), and extracellular matrix synthesis (e.g. *Omd*, *Cnmd*, *Adamts16*), whereas genes associated with matrix remodeling, mechanosensitive transcription, and immune-related signaling remained upregulated. Rather than indicating a general decline in cellular activity, this transcriptional shift reflects a transition from active growth toward tissue organization and matrix maturation.

Gene ontology analysis further substantiates this temporal adaptation (Fig. 4). During the early culture phase, enriched biological processes were dominated by environmental sensing, mechanotransduction, cell–cell communication, and responses to mechanical and metal-ion stimuli. The enrichment of neuronal-associated GO terms, including synapse assembly, cognition, learning, and memory, most likely reflects shared signaling mechanisms involved in cytoskeletal organization and intercellular communication rather than neuronal differentiation. Simultaneously, suppression of interferon- and immune-related pathways suggests that osteoblasts rapidly adapt to the curved microenvironment while limiting stress-associated responses. At day 23, GO enrichment shifted toward tissue development, cytoskeletal organization, and structural maturation. Although terms related to skeletal and striated muscle development are not osteoblast-specific, they likely represent enhanced contractility-associated and cytoskeletal remodeling programs that accompany osteoblast maturation. In parallel, reduced enrichment of migratory, chemotactic, and extracellular matrix remodeling pathways indicates progressive stabilization of cell–matrix interactions.

Collectively, these findings support a two-phase model of curvature-induced adaptation in which substrate curvature initially activates mechanosensitive signaling networks that subsequently drive extracellular matrix remodeling, tissue organization, and osteoblast maturation during prolonged culture.

### Substrate curvature primarily modulates signaling pathways involved in osteoblast differentiation

To further elucidate the influence of substrate curvature on osteoblast differentiation, we examined the expression of key osteogenic regulators and signaling pathways (Fig. 5). Despite pronounced global transcriptional changes, curvature induced only minor alterations in the expression of the master osteogenic transcription factors *Runx2, Cb@, Osx/Sp7,* and *Atf4* (Salhotra et al. 2020, Zhu et al. 2024). These findings indicate that substrate curvature does not fundamentally reprogram the core transcriptional machinery governing osteoblast lineage commitment.

However, substantial changes were observed in signaling pathways known to regulate osteoblast proliferation, differentiation, and matrix formation. Dynamic expression of *Bmp2* and *Bmp1b* suggests temporal modulation of BMP signaling during prolonged culture, while the reciprocal regulation of *Wnt8b* and *Wnt9b* indicates remodeling of WNT pathway activity as cells transition from early adaptation to later stages of differentiation. Likewise, changes in FGF family members further support the concept that curvature primarily modifies regulatory signaling networks rather than directly controlling osteogenic fate.

Notably, the increased expression of the Notch-responsive genes *Hey1, Hey2,* and *Hes3* at day 23 suggests enhanced Notch signaling during prolonged culture on curved substrates. Given the extensive crosstalk between the BMP, WNT, FGF, and Notch pathways during bone formation, these coordinated transcriptional changes likely reflect adaptive remodeling of signaling networks that regulate proliferation, matrix organization, and functional maturation rather than initiation of osteogenic differentiation itself.

These findings demonstrate that substrate curvature acts predominantly as a modulator of osteoblast signaling rather than as a direct determinant of osteogenic lineage commitment. Instead of altering the canonical osteogenic transcriptional program, curvature appears to fine-tune interconnected signaling pathways that coordinate cellular adaptation, extracellular matrix organization, and late-stage osteoblast maturation.

### Surface curvature induces remodeling of the extracellular matrix and cell–matrix adhesion complexes

The enrichment of the ECM–receptor interaction pathway indicates that substrate curvature primarily influences extracellular matrix organization and cell–matrix communication (Fig. 6). Dynamic regulation of collagens, laminins, extracellular matrix glycoproteins, and integrins demonstrates continuous remodeling of the extracellular microenvironment throughout the culture period.

Several collagen genes displayed distinct temporal expression patterns, with transient regulation of *Col26a1* and *Col9a2* contrasting with progressive upregulation of *Col9a3*, suggesting a transition from early adaptive remodeling to extracellular matrix maturation. Laminin genes exhibited comparatively modest transcriptional changes, although increased *Lamc2* expression at day 23 points to localized remodeling of basement membrane-associated matrices. Similarly, the temporal regulation of vitronectin, thrombospondin 4, and *Spp2* indicates selective remodeling of non-collagenous extracellular matrix components during prolonged culture.

Dynamic regulation of multiple integrin subunits further suggests that curvature modifies the repertoire of cell-surface adhesion receptors available for extracellular matrix sensing. As integrins are central mediators of mechanotransduction, these transcriptional changes are likely to influence downstream signaling pathways controlling osteoblast behavior and matrix organization.

The cytoskeletal expression profile (Fig. 7) indicates selective remodeling rather than global reorganization of the structural network. Differential regulation of actin isoforms, together with changes in actin-binding proteins and the marked induction of *Myh2*, suggests enhanced actomyosin contractility and dynamic adaptation of the actin cytoskeleton during prolonged culture. In contrast, the largely unchanged expression of focal adhesion and intermediate filament genes implies that these structures are predominantly regulated at the level of protein assembly, turnover, and post-translational modification rather than transcription.

Although Piezo1 and Piezo2 are recognized as key mechanosensors in osteoblasts (Zhu et al. 2024), only modest transcriptional changes were observed in response to substrate curvature (Fig. 8A). This finding is consistent with current concepts of mechanotransduction, in which Piezo channels are primarily regulated through channel activation, subcellular localization, and interactions with the cytoskeleton rather than changes in gene expression. A similar pattern was observed for septins (Fig. 8B). Despite their recently described role as membrane curvature sensors that assemble preferentially on curved membranes (Shi et al. 2023, Arnold et al. 2025), most septin family members exhibited stable transcript levels, suggesting that curvature sensing likewise depends predominantly on protein localization and higher-order filament assembly.

Collectively, these observations support a mechanistic model in which substrate curvature is primarily detected through coordinated remodeling of extracellular matrix components, integrin-mediated adhesion complexes, and the actin cytoskeleton. These structural adaptations subsequently modulate intracellular signaling pathways that regulate osteoblast maturation and extracellular matrix organization without requiring major transcriptional changes in the primary mechanosensors themselves.

## Conclusion

The present study demonstrates that substrate curvature is a potent regulator of the transcriptional response of pre-osteoblastic MC3T3-E1 cells, inducing a coordinated and highly dynamic adaptation that extends beyond the canonical osteogenic differentiation program. Although previous studies have shown that local changes in cell-membrane curvature can modulate gene expression and thus cell behaviour, these geometric changes were induced at length scales much smaller than that of a single cell (Assoian et al. 2019, Arnold et al. 2025). Till now we only had circumstantial evidence that the observed curvature sensitive responses of tissues on surfaces having small curvatures (Bidan et al. 2016, Ehrig et al. 2019) was linked to mechanosensing. The current data now demonstrates surfaces with curvatures having length scales much larger than that of a single cell, give rise to changes in transcriptome. Rather than substantially altering the expression of the master osteogenic transcription factors *Runx2, Cb@, Osx/Sp7,* and *Atf4*, curvature predominantly modulated extracellular matrix organization, cell–matrix adhesion, cytoskeletal remodeling, and interconnected BMP-, WNT-, FGF-, and Notch-dependent signaling pathways in a time-dependent manner.

Our data support a sequential model of curvature-mediated mechanoregulation. An early phase is characterized by rapid activation of mechanosensitive immediate-early genes and cellular adaptation to altered geometry, whereas prolonged culture promotes extracellular matrix remodeling, selective reorganization of the actin cytoskeleton, modulation of integrin-mediated signaling, and activation of pathways associated with osteoblast maturation and tissue organization. These findings identify surface curvature as a critical biophysical cue that regulates osteoblast function primarily by fine-tuning mechanobiological signaling networks rather than by directly reprogramming the core osteogenic transcriptional machinery.

The study further highlights the importance of incorporating surface curvature as a design parameter in next-generation bone-regenerative biomaterials and implant surfaces. Future studies integrating transcriptomic, proteomic, and functional analyses will be essential to determine how curvature-dependent changes in protein localization, cytoskeletal organization, and mechanosensor activity are translated into long-term effects on bone formation and tissue regeneration.

## Acknowledgements

The authors acknowledge the European Cooperation in Science and Technology (COST) Action EuroCurvoBioNet (no. CA22513) and the computational resources and services provided by Salzburg Collaborative Computing (SCC), funded by the Federal Ministry of Education, Science and Research (BMBWF) and the Land Salzburg.

## Authors contribution

HPO & GO: conceptualization; HPO, AL, GO: methodology; HPO, BS: investigation; AL, GO: data curation and analysis; HPO, AL, GO: visualization; HPO: writing – original draft; HPO, AL, BS, GO, JD: writing – review & editing; JD, GO: supervision

## Consent for publication

Not applicable

## Supplemental material

### Supplemental Tables

Tab. S1 Significantly differentially expressed genes.

Tab. S2 Identified genes involved in senescence.

### Supplemental Figures

**supplFigure S1.**
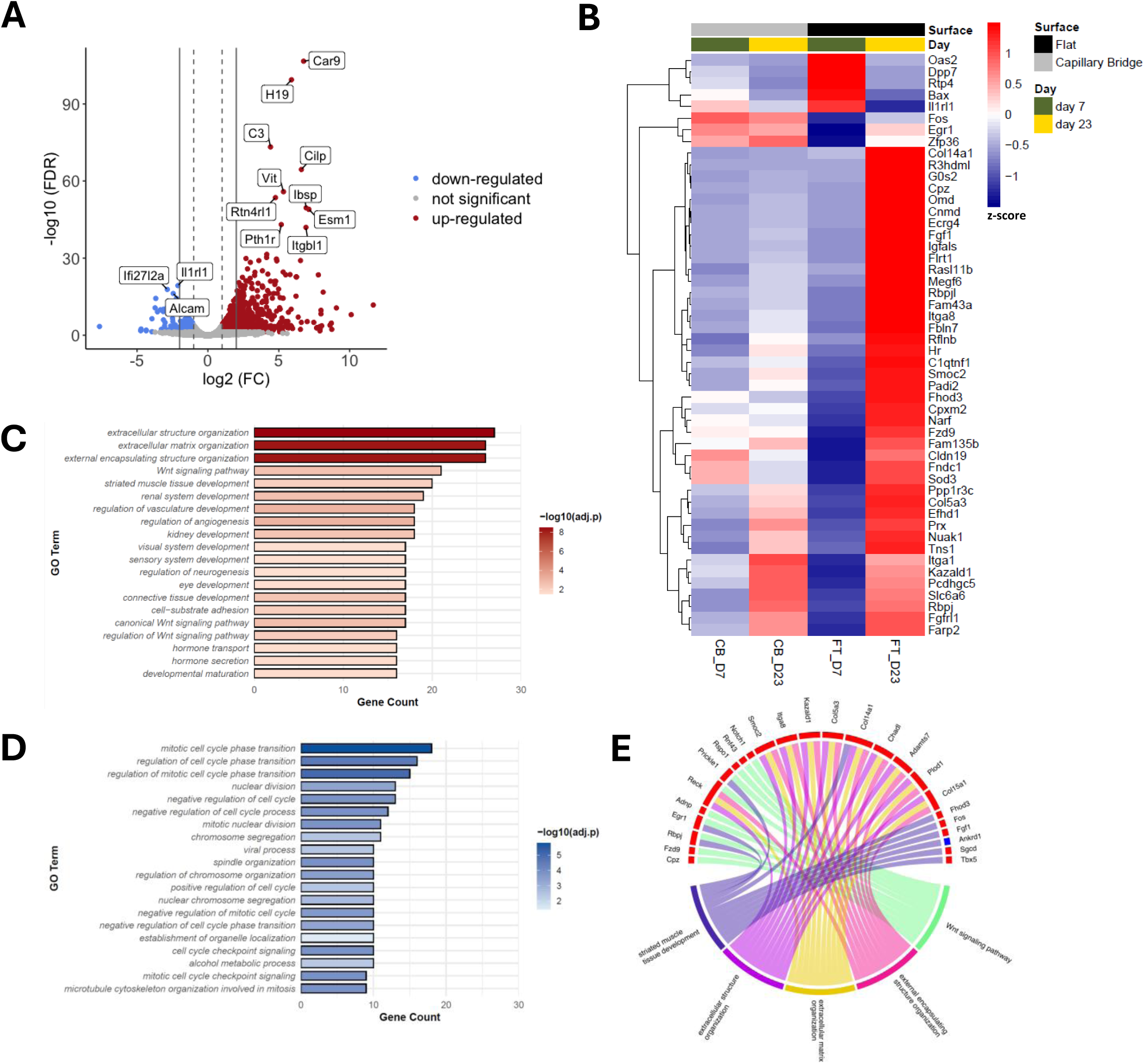
Time-dependent gene expression profiles on flat surfaces. **A.** Volcano plot visualising the up-and down-regulated genes of cells grown on flat surfaces in Petri dishes for 23 days compared to day 7. Dashed grey lines show cut-offs at log2 fold change ≥1 or ≤-1 corresponding to a doubling of values, whereas solid grey lines show a cut-off at log2 fold change ≥2 or ≤-2. The y-axis indicates the statistical significance as the log10 of the false discovery rate (FDR) corrected p-values. **B.** A heatmap shows the hierarchical clustering of the top 50 differentially expressed genes (DEGs) comparing 23-days-old against 7-days-old cells grown on flat surfaces. Z-score values of DEGs for cells cultured on flat (FT, black bar) and curved (CB, grey bar) surfaces for 7 (green bar) and 23 (yellow bar) days, respectively, are shown. Mean z-scores from replicates of the 4 conditions are given. **C.** The top 20 up-regulated biological processes in cells grown on flat surfaces. Unique DEGs after 23 days compared to 7 days-old flat-cultures were sorted to Gene Ontology categories (GO terms biological process). **D.** Top 20 down-regulated biological processes of cells grown on flat surfaces after 23 days compared to 7 days. Unique DEGs were sorted to biological process categories (Gene Ontology terms). **E.** Top 5 biological processes of all unique DEGs of cells grown on flat surfaces at day 23 compared to day 7. Individual DEGs of the same biological processes can be either up-(red) or down-regulated (blue).

